# Doberman pinschers present autoimmunity associated with functional autoantibodies: a model to study the autoimmune background of human dilated cardiomyopathy

**DOI:** 10.1101/575613

**Authors:** Gerhard Wess, Gerd Wallukat, Anna Fritscher, Niels-Peter Becker, Katrin Wenzel, Johannes Müller, Ingolf Schimke

## Abstract

**Background:** Autoimmunity associated with autoantibodies directed against the β1-adrenergic receptor (β1-AAB) is increasingly accepted as driving human dilated cardiomyopathy (DCM). Unfortunately, animal models of DCM are lacking, preventing our knowledge about β1-AAB autoimmunity in DCM from being extended and hindering the development of related treatment strategies.

**Objectives:** To introduce an animal model, we studied Doberman pinschers, which develop cardiomyopathy (DoCM), with similarities to human DCM, with regard to their β1-AAB autoimmunity.

**Methods:** Eighty-seven DP with DoCM and 31 (at enrolment) healthy controls were analyzed for β1-AAB; the receptor binding site and sensitivity to inhibition were determined. In controls who developed cardiomyopathy during the follow-up, β1-AAB were analyzed during the DoCM progress.

**Results:** Fifty-nine (67.8%) DoCM dogs and 19 (61.3%) controls were β1-AAB positive. Excluding the 9 controls who developed DoCM in the follow-up, β1-AAB positivity tended to be more pronounced in DoCM.

From the controls who developed DoCM, 8 were β1-AAB positive (p=0.044 vs. dogs remaining healthy); their β1-AAB level increased with the cardiomyopathy progress. Overall mortality and mortality exclusively due to cardiac reasons during the study period, were higher (p=0.002; p=0037) in β1-AAB positive dogs. The dogs’ β1-AAB targeted a specific epitope centralized on the second extracellular receptor and were sensitive to inhibition by drugs already successful tested for the corresponding human autoantibody.

**Conclusions:** Doberman pinschers presented β1-AAB associated autoimmunity similar to that driving the pathogenesis of human DCM. Consequently, DP could remove the lack of animal models available for studying β1-AAB autoimmunity in DCM.

## Introduction

Autoimmunity is increasingly accepted as the origin or amplifier of heart failure (1). For cardiomyopathies, preferentially for dilated cardiomyopathy (DCM) with idiopathic nature (as recently for the US re-calculated prevalence of 1 in 250-400 individuals (2)) and that secondarily to non-ischemic causes, e.g. infectious diseases such as myocarditis, several autoantibodies directed against cardiac antigens were discussed for breaking self-tolerance, leading to autoimmunity causing or supporting the disease pathogenesis (3). Among the autoantibodies, there are “classic” ones which induce immune responses, resulting in the destruction of affected tissues, whereby autoantibodies directed against contractile elements such as anti-myosin and anti-troponin autoantibodies are particularly important (4,5).

Starting in the 1970s, the classic autoantibodies were supplemented by an additional class of autoantibodies that bind to G-protein coupled receptors (GPCR-AAB). After receptor binding, GPCR-AAB, in the majority of cases, agonistically activate their related receptors in a similar way to the physiological agonists; therefore GPCR-AAB were called “functional autoantibodies”. However, mechanisms for the prevention of over-boarding receptor stimulation such as receptor down-regulation and desensitization, which are well known with physiological agonists, are lacking for GPCR-AAB. Consequently, GPCR-AAB are discussed as disease drivers as repeatedly summarized in (6–9).

With the finding of GPCR-AAB such as directed against the β1-adrenergic receptor (β1-AAB) and muscarinic receptor 2 (M2-AAB) (10,11) in patients with DCM of non-ischemic reasons, an autoimmune background of specifically associated with GPCR-AAB emerged. In contrast, patients with ischemic cardiomyopathy and healthy individuals carry GPCR-AAB only in very small amounts or these are completely absent (9).

GPCR-AAB directed against the β1-adrenerdic receptor are seen in 70–80% of patients with non-ischemic DCM (6–9).

In a distinct group of these patients, preferentially those suffering from arrhythmia, M2-AAB were additionally found with a prevalence of up to 40% (6). Among all heart specific classical and functional autoantibodies, however, the strongest pathogenicity has been demonstrated related to human DCM for β1-AAB, so that they are increasingly accepted as pathogenic drivers and treatment targets, as summarized in (6–9).

Three lines of investigation have proposed the concept of β1-AAB dependent autoimmunity in the pathogenesis of human DCM; first, experiments using myocardial cells to demonstrate the cardio-pathogenic effects of β1-AAB at the cellular and subcellular levels; second, animal experiments where rodents were immunized for β1-AAB generation to demonstrate their cardio-pathogenic effects; and third, and most impressive, clinical trials demonstrating the benefit to DCM patients when specifically their β1-AAB were removed by immunoadsorption (IA) (12,13). This treatment option is now increasingly accepted for DCM patients positive for β1-AAB. However, to overcome the problems of IA resulting from costs, logistics and patient burden, drug-associated treatment concepts for the *in vivo* neutralization of β1-AAB come increasingly to the fore (14,15). Unfortunately, to manifest and extend the knowledge about functional autoantibodies in human DCM in general and specifically of β1-AAB and even more to prove related treatment concepts in pre-clinical studies, there is a lack of animal models that are more related to human DCM than the rather artificial rodent immunization models (16,17). Although, there are also rodent models with naturally occurring DCM and transgenic mouse lines that were engineered for cardiomyopathy development. There are even mice crossed from transgenic and knockout ones which develop a “so-called” autoimmune cardiomyopathy (18,19). However, there is currently no evidence that such rodents may be suitable to eliminate the lack of models for analyzing the pathogenic role of functional autoantibody associated autoimmunity in human DCM and specifically the role of β1-AAB and M2-AAB as a driver and treatment target.

What’s more, *“rodents are phylogenetically very distant from human and some pathophysiological features of diseases and their response to pharmacological treatment may not be reliable predictors”* (18). Consequently, *“for research aimed at clinical translation, it is imperative that initial results from small rodent studies be confirmed in a large animal model that more closely resembles humans…”* (18) and there is “… a simple rule, the closer the heart or body weight of the animal to human heart and body, the more similar are the hearts” (18). Among the large animals that can be used as models for human DCM, Doberman Pinscher (DP) should be of great interest due to their frequent development of dilated cardiomyopathy (DoCM) (20), which has many similarities with human DCM (21–25).

DoCM is characterized by three stages. DP in stage one are presumed to have genetic mutations which lead to myocardial alteration on a subcellular level but the majority of cellular changes that occur is still unknown (26,27). However, affected heart mitochondrial protein expression, increased oxidative stress and evidence for apoptosis have been evidenced (28). In this stage, approximately corresponding to NYHA class 1 of human heart failure, the heart is electrically and morphologically normal (23,29). Dogs in state two (occult stage) (NYHA class 2) have either ventricular premature complexes (VPC) or a systolic dysfunction, or both, in the absence of overt clinical signs. Dogs in stage three (NYHA class 3/4) present with typical signs very similar as found in human heart failure, such as congestive heart failure (CHF), arrhythmia, syncope and exercise intolerance (20).

Here, we demonstrate for the first time that DP frequently carry β1-AAB that could act as a pathogenic driver in the pathogenesis of cardiomyopathy in a similar way to β1-AAB in human DCM. Therefore, we suggest that DP could be a suitable model for basic investigation to determine the relationship between β1-AAB-associated autoimmunity and cardiomyopathy, and even more importantly, to prove treatment concepts to counteract β1-AAB *in vivo*.

## Materials and Methods

The study was conducted in accordance with the German animal welfare law. The study protocol was approved by the “Regierung von Oberbayern”. DP were enrolled based on owner study agreement.

### Animals

Client-owned purebred DP attending the Cardiology Department of “Medizinische Kleintierklinik, Ludwig-Maximilians-Universität München” for routine check-up, cardiomyopathy diagnostics or cardiomyopathy follow-up were analyzed for β1-AAB and M2-AAB.

Based on owner study agreement a total of 118 DP (male: n=60; 50.8%, female: n=58; 49.2%) between 1 and 13 years old (median 6 years) were enrolled. To identify DoCM, Holter-ECG was performed for the detection of arrhythmia and echocardiography to evidence cardiac dysfunction. In parallel, blood was sampled for the measurement of functional autoantibodies. Based on the guidelines of the European Society of Veterinary Cardiology (30), DoCM was diagnosed for dogs with >300 VPC/24h or two subsequent examinations within a year showing between 50 and 300 VPC/24h (31) and echocardiographic indicative for cardiac dysfunction. For that purpose, the left ventricular end-systolic (ESVI) and end-diastolic volume (EDVI) were measured and indexed to body surface area based on Simpson’s method. An ESVI of >55 ml/m^2^ or/and EDVI of >95 ml^2^ were considered to be indicative of DCM.

### Measurement of autoantibodies directed against the β1-adrenergic receptor (β1-AAB) and muscarinic receptor 2 (M2-AAB)

To measure β-AAB and M2-AAB, a bioassay established by Wallukat and Wollenberger was used (11), which was modified and standardized as described in (32). In this bioassay, the chronotropic response of spontaneously beating cultured neonatal rat cardiomyocytes to the IgG prepared from the dogs’ serum was recorded (1 unit of β1-AAB activity = 1 beat/min frequency change; lower limit of detection (LLD) = 4.0 U; β1-AAB positivity = ≥ 8.0 U). Through the use of specific blockers of the β1-adrenergic (bisoprolol) and muscarinic receptor 2 (atropine), the cells’ chronotropic response can be attributed to β1-AAB or M2-AAB. For comprehensive information about sample (IgG) preparation, bioassay test setup and measurement procedure of GPCR-AAB, see (6,33).

### Localization of the receptor binding site with their specific epitope targeted by the autoantibodies directed against the β1-adrenergic receptor (β1-AAB) and muscarinic receptor 2 (M2-AAB)

To localize the extracellular binding site (loops), 50 μl of the autoantibody containing IgG preparation was pre-incubated for 30 min with 2 μl of solutions containing synthetic peptides (50 μmol/l) (Biosyntan GmbH, Berlin-Buch, Germany) which represent the first and second extracellular loops of the β1-adrenergic and muscarinic receptor 2. Then, this mixture was added to the bioassay for measurement of the autoantibodies’ chronotropic activities. To exclusively localize the extracellular binding site of β1-AAB, the bioassay was performed in the presence of 1μmol/l atropine to block the M2-AAB activity if present in the IgG preparation. To exclusively localize the target of M2-AAB, the bioassay was performed in the presence of 1μmol/l bisoprolol to block β1-AAB activity. A comparable procedure was used to map the specific epitope on the receptor loop targeted by the β1-AAB and M2-AAB. In this case, the bioassay was performed after the pre-treatment of GPCR-AAB containing IgG with an excess of synthetic peptides (Biosyntan GmbH, Berlin-Buch, Germany), which overlapped to represent the amino acid sequence of the receptor loop; first described for β1-AAB in (34). For mapping of the β1-AAB targeted epitope on the second extracellular receptor, peptides were used, as follows: P1: HWWRAE, P2: RAESDE, P3: ARRCYND, P4: PKCCDF, and P5: DFVTNR; for M2-AAB epitope mapping: P1: VRTED, P2: EDGECY, P3: CYIQFF, P4: FFSNAA P5: AAVTFG. For this, 50 μl of the GPCR-AAB-containing IgG preparation was pre-incubated for 30 min with 2 μl of solutions containing the synthetic peptides (100 μmol/l) (Biosyntan GmbH, Berlin-Buch, Germany). Then, this mixture was added to the Bioassay for GPCR-AAB measurement. In the case of finding the β1-AAB epitope, the activity was measured as described above in the presence of atropine; for M2-AAB, the bioassay was performed in the presence of bisoprolol.

### In vitro indication for the ability to neutralize Doberman pinscher autoantibodies directed against the β1-adrenergic receptor and muscarinic receptor 2 by drugs already successfully tested in animal and clinical studies for the neutralization of human autoantibodies directed against the β1-adrenergic receptor

For this purpose, the chronotropic activity of β1-AAB containing IgG from DP was monitored in the bioassay after pre-incubation of the IgG with drugs that have already been successfully tested in animal and clinical studies for β-AAB inhibition. We tested here a peptide which mimics the amino acid sequence of the second extracellular loop of the β1-adrenergic receptor (D1) and was synthesized at our request by Biosyntan GmbH, Berlin-Buch, Germany. This peptide acts comparable to the second loop mimics COR-1 which was already studied to counteract β1-AAB in patients with DCM (14). The other two substances are aptamers (15,35): the first (aptamer 110; D2) is able to neutralize only β1-AAB due to specific β1-AAB binding, which was demonstrated *in vitro* and in an animal study (36,37), while the second (BC 007; D3), as demonstrated in animal and human studies, is able to inhibit several GPCR-AAB, including β1-AAB and M2-AAB (38,39). After drug pre-incubation of β1-AAB or M2AAB containing IgG from DP with the drugs (test concentration 1 μmol/l), the mixture was added to the bioassay for measurement of the chronotropic activity of IgG.

### Statistics

Undetectable marker concentrations (<lower limit of detection, LLD) were numerically expressed as values representing one-half of the LLD. Statistical analysis was performed using the SPSS software package (SPSS Inc., Chicago, US) with Pearson chi-square test and Fisher’s exact tests for the comparison of binary variables. For the intergroup comparison of continuous data, the Kruskal-Wallis H-test combined with the Mann-Whitney U-test for post-hoc analysis for the intra-individual comparison of continuous data, and the Friedman test combined with Wilcoxon test for post-hoc analysis was employed.

For the graphical representation of continuous patient data, box plots indicate the median and interquartile range (IQR; 25th and 75th percentiles), while whiskers with ends represent the largest and smallest values inside 1.5 times the IQR, outliers (open circles) representing values between 1.5 and 3 times the IQR, and extremes (stars) placed more than 3 times the IQR.

## Results

### Basic characteristics

Among the study cohort of 118 DP as presented in Table 1, 87 (73.7%) dogs suffered from DoCM which was in age and gender composition comparable to the control group. The cardiomyopathy group consisted of dogs exclusively demonstrating arrhythmias (n=17; 19.5% - VPC/24h: median 205, min 1, max 6465; twice VPC/24 within one year: 286/173/385; EDVI: 76.45/55/91, ESVI: 40.51/22/54) indicated as the DoCM-VPC group, with exclusively echocardiographic measures outside of the reference intervals (DoCM-ECHO, n=27; 31.0% - VPC/24h: 5/0/1521; twice VPC/24 within one year: 211/0/1521; EDVI: 107.4/87/196; ESVI: 66.24/50/164) as well as those dogs presenting with arrhythmias and echocardiographic pathologies in combination (DoCM-VPC/ECHO, n=43; 49.5% - VPC/24h: 700/0/15 000; twice VPC/24 within one year: 279/124/380; ESVI: 106.8/91/160; EDVI: 67.35/42/106). The groups did not differ significantly in age. In terms of gender composition, the groups DoCM-ECHO, DoCM-VPC/ECHO and the control group are comparable, whereas in the group DoCM-ECHO female animals predominate, especially compared to the group DoCM-ECHO (p>0.05). All dogs were in the pre-clinical, occult stage of the disease. Dogs presenting with severe systemic diseases, end-stage heart failure or non-DCM cardiac diseases were excluded.

The group of dogs (n=31; 26.3% of the total number - VPC/24h: 2/0/97; twice VPC/24 within one year: 184/0/279; EDVI: 77.8/53/95; ESVI: 39.3/25/55) which did not fulfill these criteria for DoCM were defined as the primary control group (C). Related to the diagnostic criteria of DoCM used in our study, the control group was composed of healthy animals and those DP at stage 1 of DoCM. At study enrolment, 9 dogs (T0: median age 3 years; min 2, max 3 years) classified into the control group developed cardiomyopathy during the follow-up, diagnosed primarily by VPC detection (T1: median age of 7 years; min 5, max 9 years); they progressed to a diagnosis by the detection of VPCs combined with pathological echocardiography (T2: median age 9 years; min 5, max 9 years).

### Autoantibodies directed against the β1-adrenergic and muscarinic receptor 2 in Doberman pinschers at the time of enrolment

As indicated in Table 1, 78 (66.1%) of the dogs in the total study cohort presented with β1-AAB values outside of the reference range ≥8 U/min, which means that the dogs were positive for β1-AAB. Seven dogs also presented with pathological M2-AAB values, and were all also β1-AAB positive. The rest (n=40; 33.9%) presented with β1-AAB values in the reference range (< 8U/min). The dogs were sub-divided into those with DoCM and those without signs of DoCM (control group) at enrolment; however, 59 (67.8%) of the dogs with DoCM and 19 (61.3%) of the control group were positive for β1-AAB. Among the β1-AAB positive dogs, 5 of the DoCM group and 2 of the control group were also positive for M2-AAB. The remaining 28 (32.2%) in the DoCM group and 12 (38.7%) in the control group were β1-AAB negative and negative for M2-AAB. Both positivity for β1-AAB and negativity, respectively, were not significantly different between the groups. However, the median β1-AAB activity was 19.32 U/min in the DoCM group, which was slightly higher than the 16 U/min reported in the control group.

Among the control group, there were 9 dogs, 8 which were positive for β1-AAB, who developed DoCM in the follow-up. Excluding these dogs from the control group, the statistical evaluation presented a trend (p=0.097) to more β1-AAB positivity in the DoCM group compared with the control group. When reassembling the DoCM group by supplementing it with the 9 animals developing cardiomyopathy in the follow-up, the tendency (p=0.066) towards more pronounced β1-AAB positivity in the DoCM group became more marked.

Concerning the different DoCM groups, there were no significant differences related to the β1-AAB positivity (DoCM-VPC: n=12, 70.6%; DoCM-Echo: n=15, 55.6%; DoCM-VPC /Echo: n=32, 74.4%).

**Table 1.**
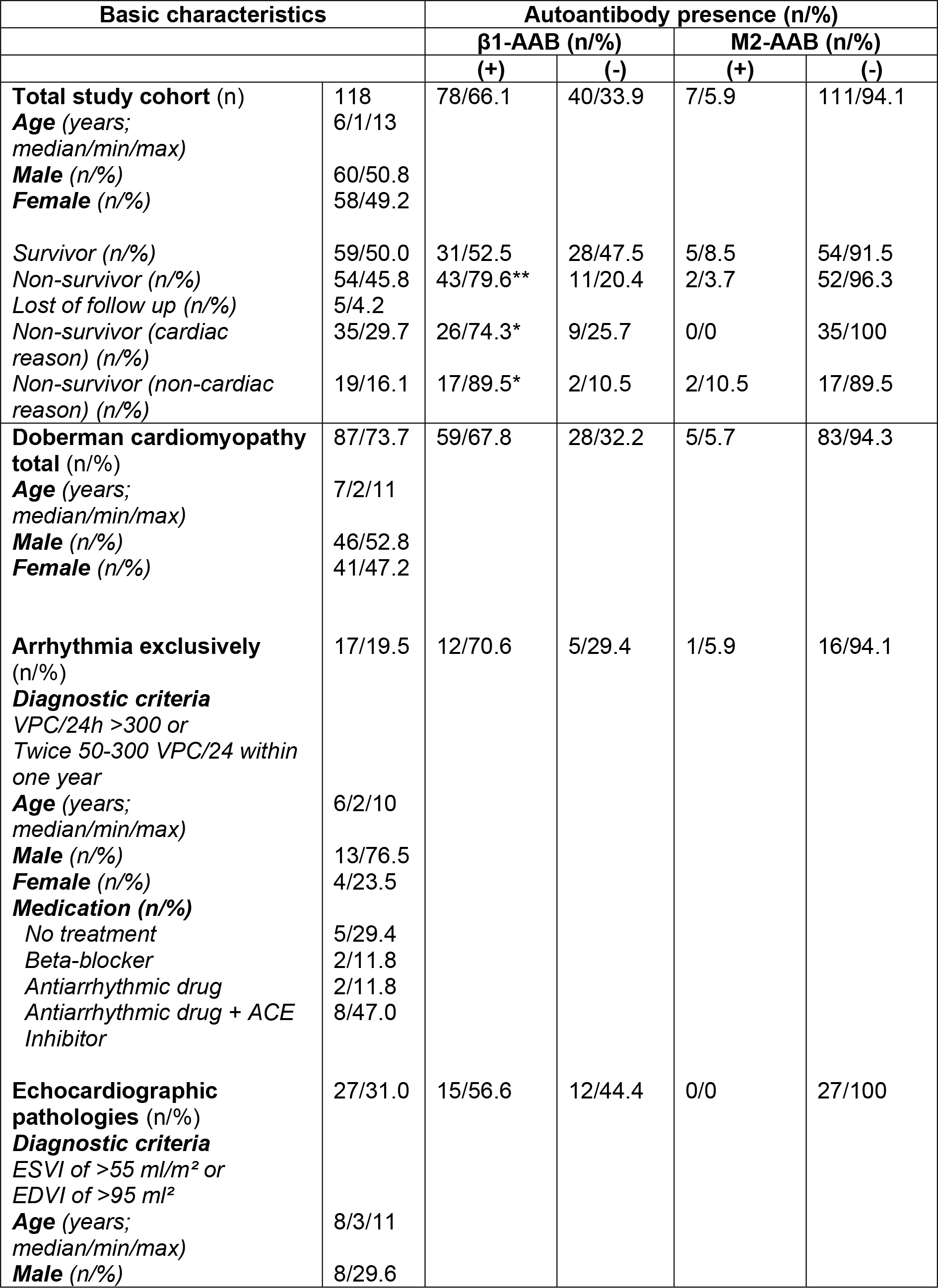
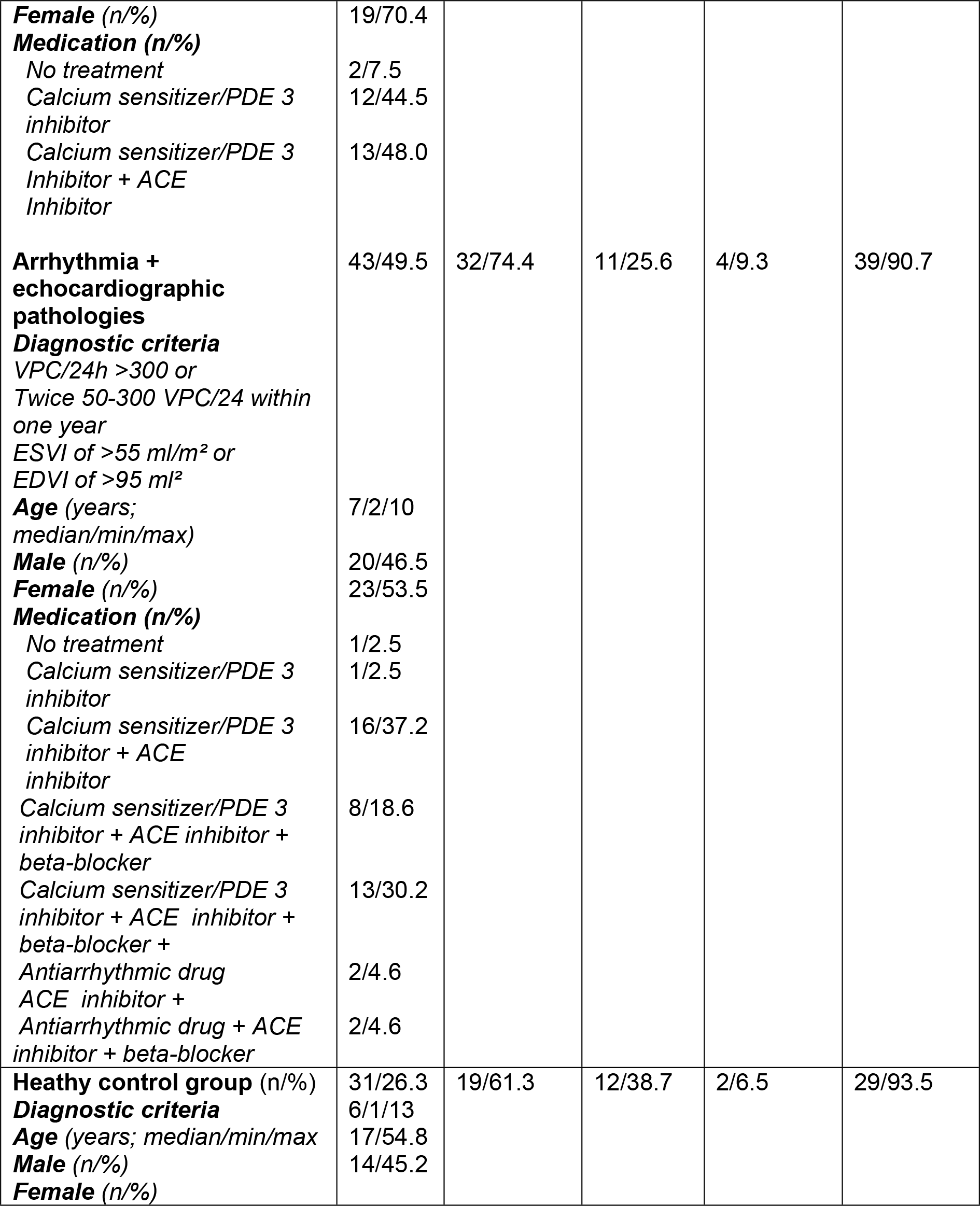
Basic characteristics and serum activities of autoantibodies directed against the β1-adrenergic and muscarinic receptor 2 (* p<0.05; ** p<0.01)

### Follow-up of autoantibodies directed against the β1-adrenergic receptor in primary healthy dogs who progress to cardiomyopathy

Among the 9 dogs which were healthy at the time of enrolment (DoCM T0) but progressed to DoCM, one of these animals was negative for β1-AAB at enrolment and remained negative despite progressing to DoCM diagnosed by VPC (DoCM T1) and later than by VPC combined with pathologic echocardiography (DoCM T2). The other 8 dogs were β1-AAB positive at enrolment. In parallel to the cardiomyopathy development, the β1-AAB levels increased from T0 to T1 in all DP (p<0.05) and in β1-AAB positive DP at study enrolment (p<0.02), from T1 to T2 in all DP (n.s) and in β1-AAB positive DP at study enrolment (p<0.02), as well as from T0 to T2 in all DP (p<0.05), and in β1-AAB positive DP at study enrolment (p<0.02). A further rise in β1-AAB from T1 to T2 was demonstrated for seven DP. In the one dog that showed a decrease in the β1-AAB level from T1 to T2, the β1-AAB level remained clearly in the pathological range (Figure 1).

In the comparison of dogs who were free of DoCM for the whole study period with those without the signs of DoCM at enrolment but who progressed to DoCM, a significantly higher proportion of β1-AAB positivity (p=0.044) was calculated for the last animals.

**Figure 1.**
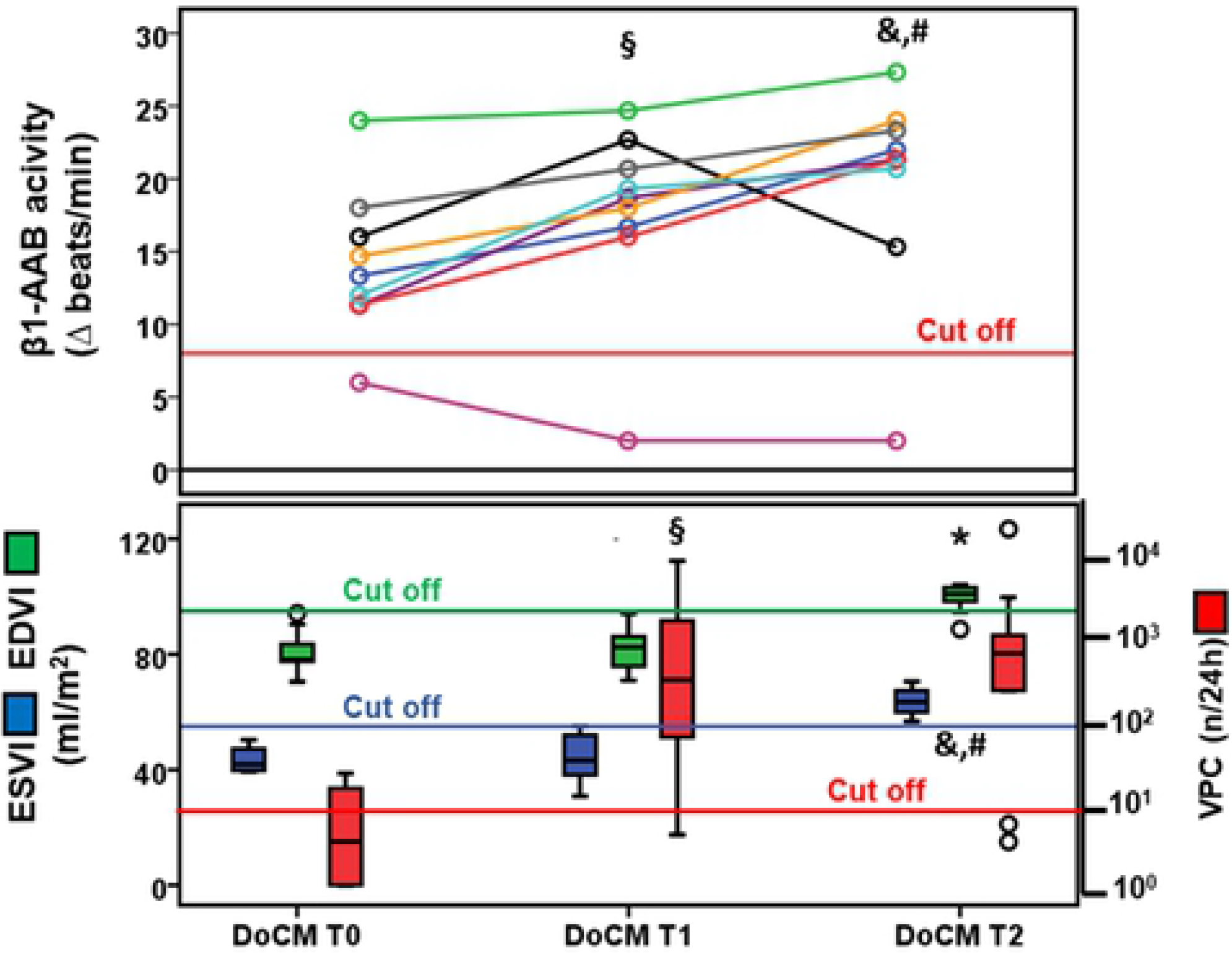
(A) Activity of autoantibodies directed against the β1-adrenergic receptor and (B) left ventricular end-systolic (ESVI), and end-diastolic volume (EDVI) indexed to body surface area and ventricular premature contractions per 24 hours (VPC/24h) in primarily healthy Doberman pinschers (DP) (n=9) during the development of severe cardiomyopathy. (DoCM T0 = healthy; DoCM T1 = cardiomyopathy indicated by arrhythmia; DoCM T2 = cardiomyopathy indicated by arrhythmia combined with pathological echocardiography); **(A)** § T1 vs. T0: p<0.05 in all DP, p<0.02 in β1-AAB positive DP at study enrolment; & T2 vs. T1: p<0 02 in β1-AAB positive DP at study enrolment; # T2 vs. T0: p<0.05 in all DP, p<0.02 in β1-AAB positive DP at study enrolment; **(B)** § T1 vs. T0: p<0.01 (VPC724h), & T2 vs. T1: p<0.02 (VPC/24h), p<0.01 (EDVI, ESVI), # T2 vs. T0: p<0.02 (VPC724h, EDVI), p<0.01 (ESVI).

### Mortality of Doberman pinschers related to autoantibodies directed against the β1-adrnergic receptor

Related to the 118 dogs enrolled, 59 (50%) survived the study period, 35 (29.7%) died due to cardiac reason such as sudden death (n=30; 85.7%) or heart failure (n=5; 14.3%) and 19 (16.1%) died from non-cardiac reasons (Table 1). The median survival time of dogs related to the time of enrolment was 1 year (min 0, max 9 years). Five (4.2%) dogs were lost to follow-up. Of the surviving dogs, 28 (47.5%) were β1-AAB negative at study enrolment and 31 (52.5%) were β1-AAB positive. In contrast, only 11 (20.4%) of the non-survivors were β1-AAB negative while 43 (79.6%) were positive for β1-AAB, which documents a significantly higher prevalence (p<0.01; odd ratio 3.61 (1.57-8.33) of β1-AAB positivity in the non-survivors.

The increased prevalence of β1-AAB in the non-survivors concerned the dogs that specifically died due to cardiac reasons (n=9; 25.7% β1-AAB negative vs. n=26 (74.3%) β1-AAB positive; p<0.05; odds ratio 2.61 (1.05-6.51) but also those died due to non-cardiac reasons (p<0.05; odds ratio 7.93 (1.68-37.49). With respect to M2-AAB, 54 (91.5%) of the surviving dogs were negative at study enrolment and 5 (5.7%) were positive. However, M2-AAB positivity did significantly increase the risk for death.

### Characteristic features of autoantibodies directed against β1-adrenergic and muscarinic receptor 2 present in Doberman pinschers

As exemplarily demonstrated for β1-AAB in Figures 2 and 3, both β1-AAB and M2-AAB found in DP targeted the second extracellular loops of the related receptors. With respect to the epitope on the second extracellular loops which were targeted, the β1-AAB epitope is located more centrally and contains 3 cysteine residues while the M2-AAB epitope is located closer to the N-terminus and is only flanked by 1 cysteine.

**Figure 2.**
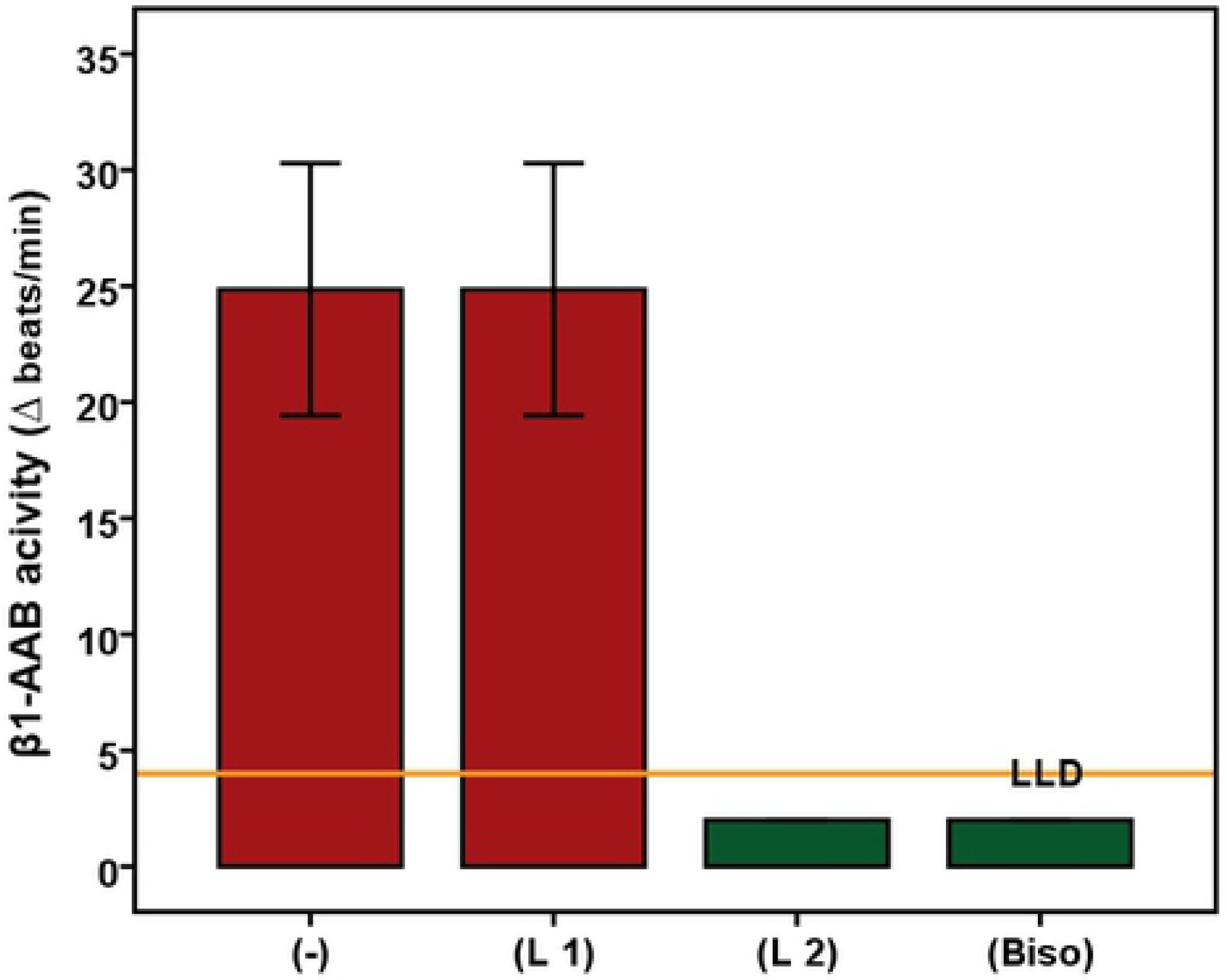
Autoantibodies directed against the β1-adrenergic receptor (β1-AAB) of Doberman pinschers (n=6) target the second extracellular receptor loop. Using the bioassay of spontaneously beating cultured neonatal rat cardiomyocytes, the chronotropic activities of the Doberman pinschers’ β1-AAB is demonstrated by the absence or presence of the peptides (L1 = first loop; L2 = second) competing with the first and second extracellular receptor loops. The control experiment was performed in the presence of bisoprolol (BISO). Values below the low limit of detection (LLD) were displayed as half range values. LLD = 4 beats/min; cut off (separating healthy from disease subjects) = 8 beats per/min.

**Figure 3.**
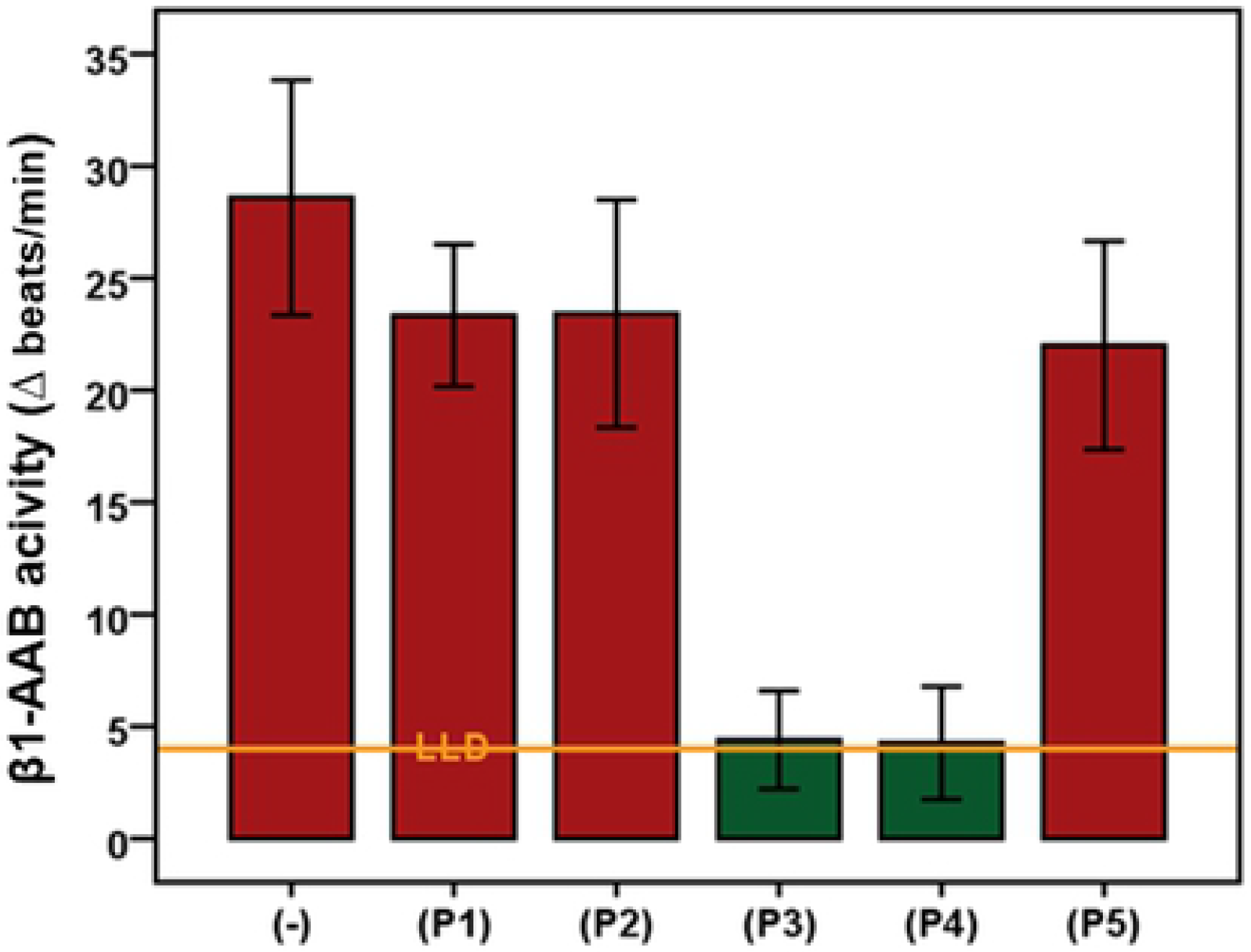
Mapping of the second extracellular loop of the β1-adrenergic receptor for epitope localization targeted by the related autoantibodies (β1-AAB) of Doberman pinschers. Using the bioassay of spontaneously beating cultured neonatal rat cardiomyocytes, the β1-AAB (n = 5) were measured in the absence or presence of competing peptides that overlapped to represent the second extracellular receptor (P1: HWWRAE, P2: RAESDE, P3: ARRCYND, P4: PKCCDF, and P5: DFVTNR). Values below the low limit of detection (LLD) were displayed as half range values. LLD = − 4 beats/min; cut off (separating healthy from disease subjects) = − 8 beats per/min.

Based on bioassay measurements, Figure 4 demonstrates that all three drugs which were successful for the neutralization of β1-AAB in human with DCM were also able to neutralize DP β1-AAB. Compared with the original β1-AAB containing DP IgG, the same IgG pre-incubated with the drugs did not present any chronotropic activity on spontaneously beating neonatal cardiomyocytes.

**Figure 4.**
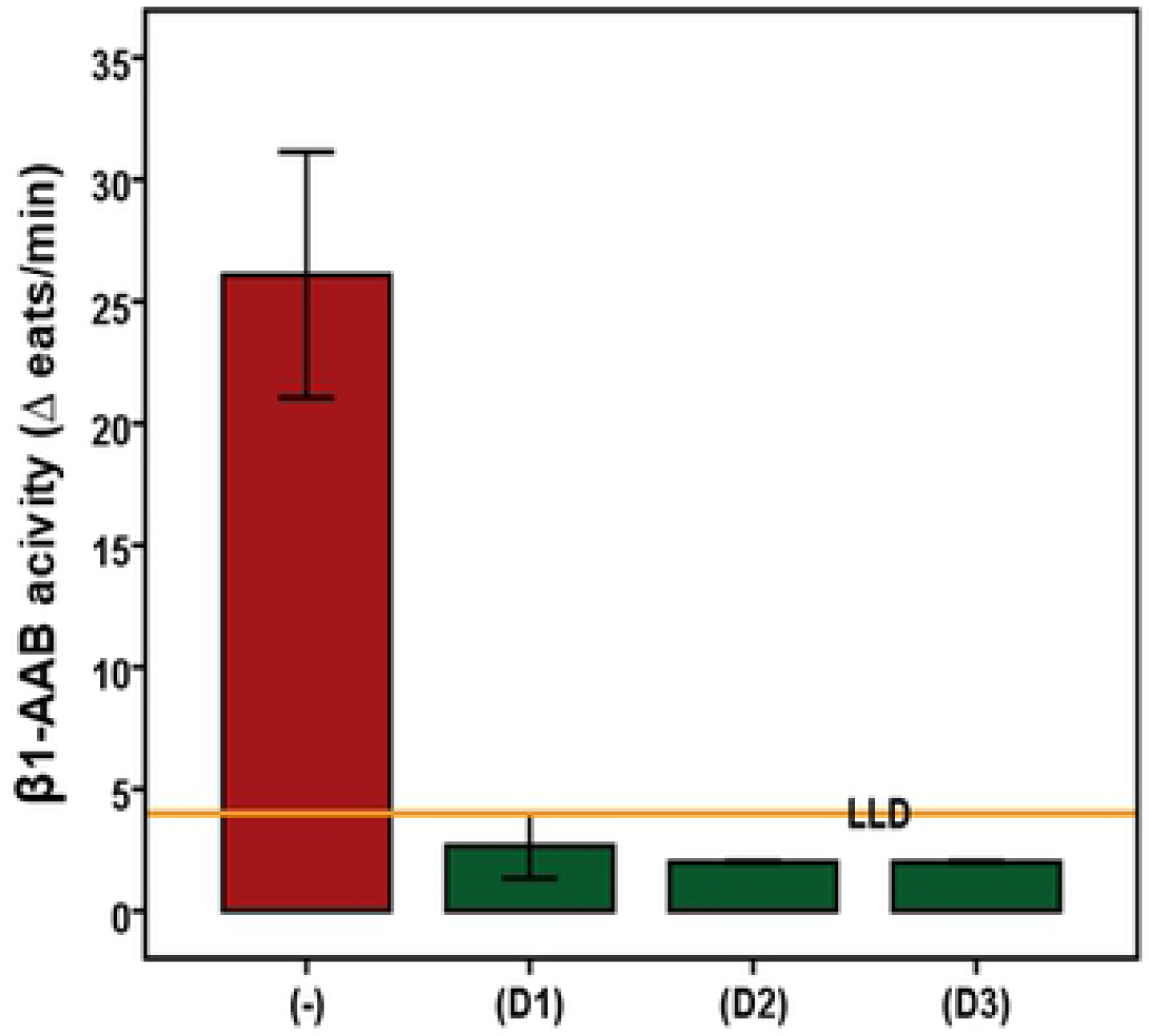
In vitro indication for the ability to neutralize Doberman pinscher autoantibodies directed against the β1-adrenergic receptor by drugs documented to neutralize human autoantibodies directed against the β1-adrenergic receptor. Using the bioassay of spontaneously beating cultured neonatal rat cardiomyocytes, the β1-AAB were measured in the absence (control n=8) or presence of the drugs (D1 = second loop peptide (n=4), D2 = aptamer 110 (n=2), D3 = aptamer BC 007 (n=4)). Values below the low limit of detection (LLD) were displayed as half range values. LLD = − 4 beats/min; cut off (separating healthy from disease subjects) = − 8 beats per/min.

## Discussion

Dilated cardiomyopathy with a disease cumulative prevalence of 58% (20,30,40,41) is the most common form of cardiomyopathy in Doberman pinschers which “… closely resembles the human form of the disease” (24) and therefore repeatedly suggested for modeling human DCM (18,30). In the final state, as mentioned already in above, DP with DoCM present with typical signs such as congestive heart failure (CHF), arrhythmia, syncope and exercise intolerance very similar as found in human heart failure. From anatomical and morphological points of view, left ventricular chamber dilatation and fibrotic cardiac rearrangement were seen (20).

The majority of dogs (93.5%) do not survive 2 years after being diagnosed with DoCM (42), which is in agreement with our findings. Despite optimal treatment, the survival of the dogs is about 130 days (median) after entering the overt stage (43).

Almost 30 years ago, Smucker et al. (25) suggested that DP should be used as a model for human DCM. Subsequently, DP with dilated cardiomyopathy were announced as *“…remain(ing) (an) untapped resources to investigate both mechanisms of arrhythmias and pharmacodynamics of anti-arrhythmics”* (21).

However, DP as a DCM model did not gain widespread acceptance in basic research and also not in pre-clinical studies for the testing of human drugs.

For the pathogenesis of DoCM, a genetic background is discussed and a autosomal dominant inheritance was proposed (41) but it has been stated (30) that *“… absence of a (specific) genetic mutation … associated with … DoCM … does not ensure the dog will never … develop DCM … (and) … identification of a genetic mutation does not guarantee the dog will … develop DCM”*. Comparable to the genetic background discussed for DoCM, genetic reasons are also assumed to be prominent in human DCM (44). A familial disease history was found in 25% of the human DCM patients (45) detected by a highly diverse genetic background with several gene mutations that, nevertheless, produces a relatively unique DCM phenotype (2).

For this complexity in the pathogenesis of DoCM and human DCM, consequently, further causes such as the phenomenon of autoimmunity against the heart must be considered whereby it is currently beyond any doubt that autoimmunity is in tight relation to the individuals’ genetic backgrounds. (46).Today, it is increasingly accepted that autoimmunity is as an important pathogenic driver of human DCM (47) and functional autoantibodies such as β1-AAB and M2-AAB diseases came to the fore (6–9). Finding a comparable autoimmunity in DoCM, additionally to all the other similarities of DoCM with human DCM summarized (21–25), would Doberman pinchers predestinate as model to investigate the autoimmune background of human DCM in general and specifically treatment strategies directed to the functional autoantibodies. We present here for the first time data which indicate that the autoimmune background associated preferentially with β1-AAB and discussed as driver of human DCM also probably drive the pathogenesis of DoCM.

In our study cohort, DP with DoCM showed a prevalence of β1-AAB of about 70%, comparable as is in human DCM (6). Furthermore, non-surviving dogs were significantly more often positive for β1-AAB than survivors, which clearly agrees the findings with patients with human DCM studies (48–50).

Interestingly, the dogs of our control group also carried β1-AAB (about 60%). Whether the β-AAB positive dogs are those dogs being genetically compromised for DoCM and therefore in stage one of DoCM remains speculative as it remains whether these dogs are those who progress to the clinically overt DoCM. However, 60 % of β1-AAB positivity in the control group corresponds to the documented DoCM prevalence (20). Additionally, there was a significantly higher frequency of β1-AAB positivity in the healthy dogs who developed DoCM during the study period which could support the assumption of β1-AAB dependent driving to clinically relevant DoCM. Related to humans, we discussed the role of β1-AAB autoimmunity in progressing to cardiomyopathy for Chagas’ patients, where 30% of asymptomatic patients were positive for β1-AAB and, based on epidemiologic data, nearly 30% of asymptomatic Chagas’ patients also progress to Chagas’ cardiomyopathy (32). Taking all of this together, we assume a prominent driving role for β1-AAB associated autoimmunity in the pathogenesis of DoCM, as is increasingly being accepted for human DCM. The resembled role of β1-AAB autoimmunity in the pathogenesis of DoCM and human DCM, for humans again deduced from Chagas’ patient data, was also supported by the increase in the β1-AAB level from healthy or asymptomatic subjects to those suffering from mild to severe cardiomyopathy, which was seen in both Doberman pinschers and Chagas’ patients (32). Consequently, measurement of β1-AAB could be potentially used for monitoring and prognosis of DoCM and Chagas’ cariomyopathy. Unfortunately, corresponding longitudinal studies focused directly on patients developing DCM are still lacking, but we hope that bio-banking concepts will facilitate the access of such data in the near future.

If we take a look at the characteristics of dog and human β1-AAB, some further analogies were obvious. β1-AAB of Doberman pinschers target the second extracellular receptor loop, where there is an epitope localized centrally and containing a cysteine between amino acids 193 and 204 which is comparable with the epitope targeted by β1-AAB found in DCM patients (34,51); however, it must be stated that there are additional β1-AAB in human DCM which target the first extracellular receptor loop (34) which have not been seen in Doberman pinschers. However, β1-AAB directed against the second extracellular receptor loop were sometimes (52) but not always (34) accused of being the determining cause of DCM.

As for β1-AAB of DCM patients described in (14,37,39), the β1-AAB activity of Doberman pinschers could be inhibited by peptides which mimic the second extracellular receptor loop or by aptamers binding the autoantibodies. In our view, this is a further indicator that the β1-AAB associated autoimmunity in Doberman pinschers and human is closely related. Although M2-AAB were found with a clearly lower prevalence in DoCM than published for human DCM, the characteristics are comparable. M2-AAB targeted the second extracellular receptor loop which was also published for M2-AAB of human cardiomyopathies, such as DCM and Chagas’ cardiomyopathy (53,54). The specific epitope is located closer to the N-terminus, between amino acids 169 and 177; the same region was also demonstrated for M2-AAB of patients with DCM (unpublished data) or Chagas’ cardiomyopathy (54). Related to their specificity for β1-AAB inhibition, the second loop peptide (D1) and aptamer 110 (D2) did not inhibit M2-AAB, but the aptamer BC 007 (D3), the so-called “broad-band neutralizer” of GPCR-AAB, inhibited the Doberman pinscher M2-AAB as seen for M2-AAB from DCM patients (39,55).

Consequently, we suggest, to take the information about functional autoantibody associated autoimmunity in DoCM, together with all the other similarities of DoCM with human DCM as summarized in (21–25), to re-activate Doberman pinschers as a model of human DCM, specifically for basic investigation of the functional autoantibody associated autoimmunity in human DCM and still more importantly for pre-clinical studies in the development of treatment strategies directed against functional autoantibody associated autoimmunity.

From our point of view, this is all the more important since, firstly, none of the small animal models used for the modelling of human DCM to date develop functional autoantibodies and, secondly, the functional autoantibodies found in the immunization models seem, based on ELISA experiments (56) to differ from the human autoantibodies in quality and quantity.

We agree that a large animal model such as the DP is a priori cost-intensive and there are strong requirements based on “World Medical Statement on Animal Use in Biomedical Research” to respect the welfare of animals in general and specifically of large animals such as DP used for research. However, a study design such as used for the present study with enrolment of client-owned purebred DP attending a veterinary-medical institution for routine check-up, disease diagnostics or follow-up would strongly minimize the cost and guarantee the DP’s welfare. In addition, the DP of our study came from different breeding populations throughout Europe and therefore has a greater genetic diversity than the DP from one litter or only a few and can therefore better reflect the pathogenic situation in human DCM with its already mentioned very different genetic background.

Last but not least, we have learned that dog owners have a great willingness to participate with their dogs in studies that test new treatment options, especially if it is expected that the treatments can also be beneficial to their dog; always provided that a study design is chosen that guarantees a minimal physiological and psychological impairment of the animals, which was ensured in our study by non-invasive heart examination and only blood analysis.

## Study limitations

Based on the diagnostic criteria for DoCM used in our study, the control group consisted of healthy animals and DP at stage 1 of DoCM. It is assumed that the dogs at this stage exhibit genetic mutations that lead to myocardial alteration at the subcellular level without becoming electrically or echocardiographically visible. We suspected that the β1-AAB-positive DP of the control group were those at level 1 of the DoCM. In the future, a detailed characterization of the DP with respect to a genetic predisposition is necessary to verify this speculation.

## Conclusions

Doberman pinschers with cardiomyopathy presented with typical signs for autoimmunity, preferentially with autoimmunity associated with autoantibodies directed against G-protein coupled receptors, which is closely related to the autoimmunity found in patients with DCM. This, together with the higher prevalence of cardiomyopathy in Doberman pinschers, the fast disease progression and reaching the primary end point of death within around 4 months of entering the severe stage of cardiomyopathy, presents an excellent basis to re-activate Doberman pinschers as a model to study the basics of human DCM. This is specifically the case for GPCR-AAB associated autoimmunity as a disease cause and as a model for pre-clinical studies in drug development aimed at counteracting this form of autoimmunity and consequently DCM.

